# Mechanical memory in cells emerges from mechanotransduction with transcriptional feedback and epigenetic plasticity

**DOI:** 10.1101/2020.03.20.000802

**Authors:** Jairaj Mathur, Vivek B. Shenoy, Amit Pathak

**Affiliations:** Department of Mechanical Engineering & Materials Science, Washington University, St. Louis; Department of Materials Science and Engineering, University of Pennsylvania, Philadelphia PA; Center for Engineering MechanoBiology, University of Pennsylvania, Philadelphia PA

**Keywords:** mechanical memory, mechanotransduction, transcription, epigenetic plasticity, mechanobiology

## Abstract

Emerging evidence shows that cells are able to sense and store a memory of their past mechanical environment. Since existing mechanotransduction models are based on adhesion and cytoskeletal dynamics that occurs over seconds and minutes, they do not capture memory observed over days or weeks. We postulate that transcriptional activity and epigenetic plasticity, upstream of adhesion-based signaling, need to be invoked to explain long-term mechanical memory. Here, we present a theory for mechanical memory in cells governed by three key components. First, cells on a stiff matrix are primed by a transcriptional reinforcement of cytoskeletal signaling. Second, longer stiff-priming progressively produces more memory-regulating factors and reduces epigenetic plasticity. Third, when stiff-primed cells move to soft matrix, the reduced epigenetic plasticity blocks new transcription required for cellular adaptation to the new matrix. This stalled transcriptional state gives rise to memory. We validate this model against previous experimental findings of memory storage and decay in epithelial cell migration and stem cell differentiation. We also predict wide-ranging memory responses for different cell types of varying protein kinetics and priming conditions. This theoretical framework for mechanical memory expands the timescales of mechanotransduction captured by conventional models by integrating cytoskeletal signaling with transcriptional activity and epigenetic plasticity. Our model predictions explain mechanical memory and propose new experiments to test spatiotemporal regulation of cellular memory in diverse contexts ranging from cell differentiation to migration and growth.

## INTRODUCTION

As living cells engage their receptors with protein ligands present in their extracellular matrix (ECM), the resulting cell-ECM adhesions assemble differently according varying mechanotransduction phenomena, including cell stretch, elevated hydrostatic pressure, fluid shear stress, osmotic forces, matrix confinement, and substrate stiffness (1–7). Specifically, on stiffer matrices, adhesion bonds support higher forces with greater clustering of focal adhesion proteins (8, 9), which ensues complex bio-chemo-mechanical processes, stress fiber contractility (10), and signaling pathways collectively known as mechanotransduction (11, 12). Over the past two decades, the machinery of mechanotransduction has formed the foundational principles for the field of mechanobiology. Several mathematical models have described and revealed in great detail how cells incorporate various biochemical signaling events mediated by forces in focal adhesions and stress fibers to generate varying degrees of mechanoactivation, which in turn governs basic cellular responses such as migration and proliferation. For example, a motor-clutch description of forces and adhesions captures the known biphasic dependence of cell migration and spreading on ECM stiffness (13–15). Phenomenological integration of adhesions, forces, and cellular polarity predicts varying regimes of biphasic responses in matrices that differ in stiffness and confinement (16, 17). In fibrous environments, adhesion dynamics tends to match the timescales of ECM viscoelastic response (18). These and numerous other models have provided fundamental insights into cell-matrix interaction and mechanotransduction (19–21). However, current understanding of mechanotransduction is limited to a given timepoint and cells sensing their immediate microenvironment, without taking into consideration the effect of subcellular processes that take place at longer timescales of days and weeks.

In recent years, emerging evidence has shown that cells can be primed or dosed by a given matrix stiffness and they continue to show signatures of their past priming even after they move to a new matrix via a stored mechanical memory (22–24). Priming of stem cells on stiff matrix localizes YAP to the nucleus, activates RUNX2, and reduces stem cell plasticity, with a bias towards the osteogenic lineage. Such ‘stiff-priming’ of stem cells over longer duration causes more stable memory response, lasting 3-10 days, measured in terms of YAP activation and osteogenic markers (23). Separately, miRNA-21 transcription and downstream MRTF signaling have been implicated in long-term memory in stem cells over several weeks (22). Based on prior findings of mechanosensitive stem cell differentiation (23, 25), a recent model (26) accounted for YAP/TAZ kinetics in a continuous range of stiffness values and predicted that different mechanical memory regions exist for four specific stem cell lineages. However, this model does not capture how memory responses degrade as substrate stiffness changes over time. Since memory-regulating genes for cell types and situations other than mesenchymal stem cell fate decisions remain largely unknown, there is a need to understand how cytoskeletal signaling, transcriptional activity and epigenetic modifications combine to generate broad memory responses for biological processes beyond stem cell differentiation. In the context of cell migration, we have recently shown that epithelial cell sheets primed on a stiff matrix for 3 days store mechanical memory through nuclear YAP localization, which continues to enhance cell migration through enhanced pMLC expression and focal adhesion formation on soft matrix for 2-3 days (24). Priming less than 3 days substantially reduces memory, which suggests that memory processes operate progressively over time.

The subcellular processes that potentially govern memory over days/weeks occur at much longer timescales than the conventional processes in cellular mechanotransduction, such as adhesion formation and stress fiber contractility, that operate and turnover in minutes/hours. We propose that mechanosensitive transcriptional activity upstream of cytoskeletal signaling, such as YAP, miR21 and MRTF (4, 22–24), is responsible for producing stiffness-specific proteins, such as pMLC, vimentin and αSMA (22, 24). Previously, Li *et al.* proposed that stiff-priming of cells progressively fills a so-called “reservoir” of memory through transcriptional activity and downstream signaling over time (22). Once the reservoir is filled with the levels of miR21, YAP/TAZ and MRTF, cells are ‘stiff-primed’ and they continue to show fibrotic response until these memory regulators are dissipated. However, it is not clear why the cells do not commence new transcription and protein signaling as soon as they move to the soft matrix. Additionally, the reservoir model does not capture how the production and degradation kinetics of memory regulators, such as YAP, MRTF, and miR21, is coupled with the matrix-sensing cytoskeletal processes that ultimately govern cellular outcomes of memory-dependent migration or differentiation (22–24).

In this manuscript, we address gaps in the current understanding of mechanotransduction and memory regulation over longer time scales of days to weeks through a novel conceptual and mathematical model, which connects cytoskeletal mechano-sensing to nuclear mechanoactivation and epigenetic plasticity. Beyond the classic notion of cytoskeletal response to matrix stiffness, recent studies have shown that nuclear morphology and chromatin accessibility also depend on matrix stiffness (27–31). Thus, it is likely that the matrix-dependent epigenetic changes feedback to cytoskeletal signaling. Since the epigenetic landscape fundamentally regulates transcription factor (TF) binding and gene expression (32–34), the connection between stiffness-sensitive cytoskeletal signaling and epigenetic programming is a critical missing piece in the theoretical framework of mechanotransduction. Indeed, when stem cells are stiff-primed and moved to a soft matrix, an epigenetic modification factor HAT1 is persistently expressed and histone deacetylation enzymes (HDACs) remain low (35), both of which enhance chromatin resistance. Based on this work and related experimental findings of priming-dependent cellular response (22–24, 35), we propose that mechanical priming of cells requires the production of memory-regulating factors, such as epigenetic modifiers (e.g., HAT1, HDACs) and nuclear structure regulators (e.g., Lamin A/C). These memory factors reduce epigenetic plasticity, defined as the ability of cells to alter their chromatin configuration and expose TF binding sites.

In the context of memory, when a stiff-primed cell is moved to a soft matrix, the reduced epigenetic plasticity blocks the transcription of new genes required for adaptation to the new environment. As a result, the prior stiff-primed cellular state persists until the memory factors are degraded and the epigenetic plasticity is restored. We integrate these key processes corresponding to transcriptional activity, memory factor production, downstream signaling, epigenetic plasticity, and combined mechanoactivation from nuclear and cytoskeletal contribution, all operating at varying kinetics, in a new expanded mathematical framework for mechanotransduction. Our calculations validate existing experimental findings on mechanical memory and predict new outcomes that can potentially reveal the underlying mechanisms of memory regulation.

## THE MODEL

### Combining cytoskeletal and nuclear contributions

We define a dimensionless parameter called “cellular mechanoactivation” (*ϕ*) that captures all classic signals that go up on stiff substrates and down on the soft ones, such as nuclear YAP localization (36), actin-myosin forces (37), focal adhesion formation (11), and mesenchymal markers (38, 39). This broad definition allows us to focus on the newly introduced differential kinetics of nuclear and cytoskeletal signaling. Next, we postulate that cellular mechanoactivation *ϕ* receives contributions from both cytoskeletal and nuclear mechanotransduction, *ϕ_c_* and *ϕ_n_*, respectively (Fig. 1). We model a coarse-grained net representation of the *ϕ* machinery as:

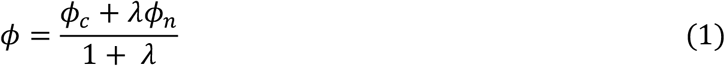

**Figure 1:**
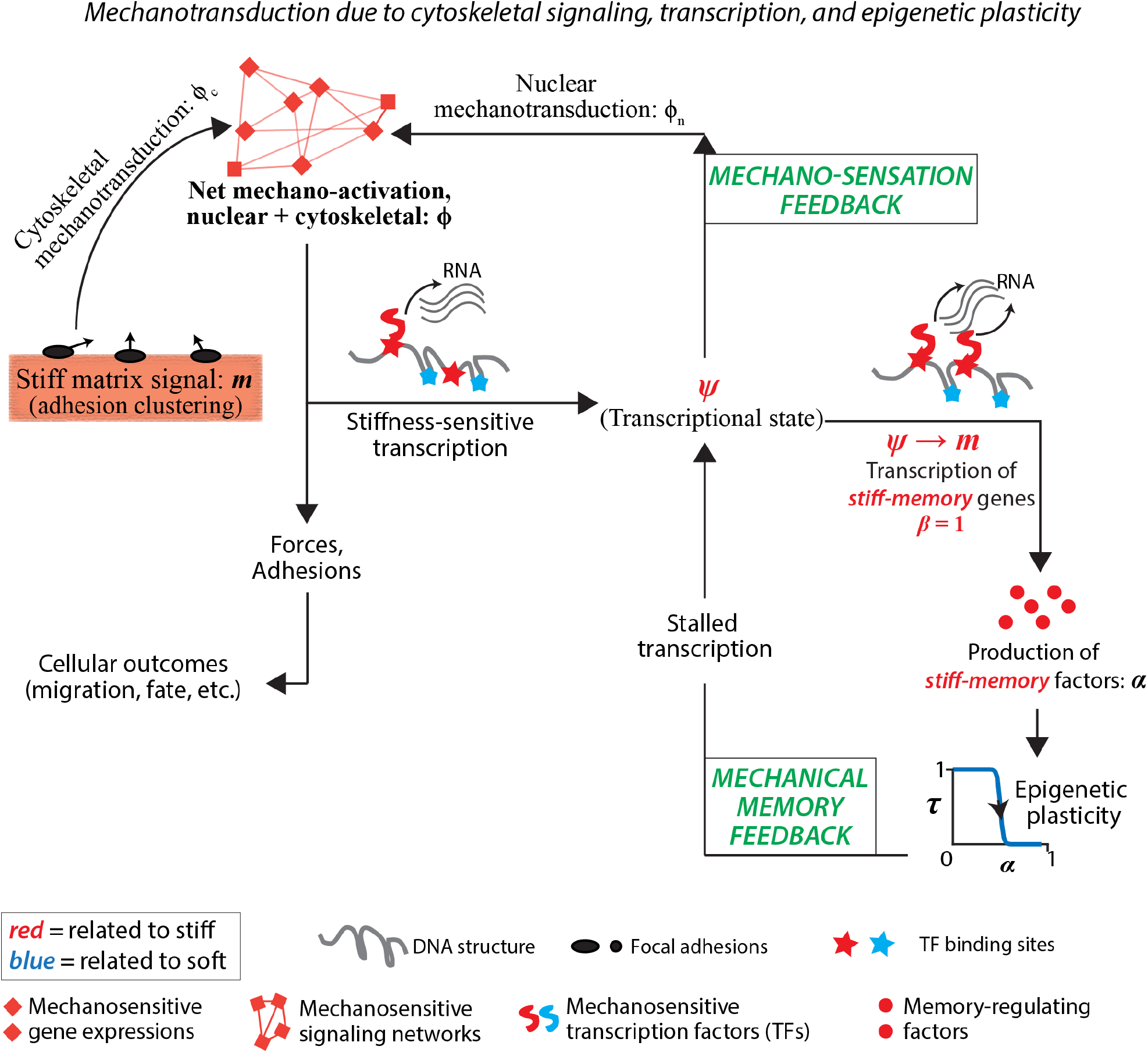
A model for mechanotransduction through feedback from transcriptional activity and epigenetic plasticity. Schematic describing key events in the process of mechanotransduction as cells sense and respond to stiff matrices. Cells sense matrix stiffness through adhesions and contribute to cytoskeletal mechanotransduction (*ϕ_c_*). Cellular mechanoactivation allows stiffness-sensitive epigenetic modifications, which expose binding sites for stiffness-specific transcription factors. As a result, cells achieve mechanosensitive transcription (*ψ*) and gene expressions that dictate nuclear mechanotransduction (*ϕ_n_*). This interdependence of matrix signal (*m*), cytoskeletal signaling, transcription, and nuclear mechanotransduction establishes a ‘mechano-sensation feedback’. When the transcriptional state fully adapts to the matrix signal (*ψ* → *m*), we propose that transcription towards memory-dependent gene expression commences (*β* = 1), producing factors like epigenetic modification factors, that establish a “stiff-memory” and reduce epigenetic plasticity (*τ*). As a result, the genome is locked in its current state and the transcription required for adaptation to new matrix stiffness is stalled, which gives rise to a ‘mechanical memory feedback’ of the stiff matrix. Stalled transcription leads to continued upregulation of the mechanosensation feedback, resulting in upregulated mechanoactivation.

Here, *ϕ_n_* represents the nuclear contribution towards cytoskeletal processes due to mechanosensitive mRNA production via stiffness-sensitive transcriptional activity, such as nuclear YAP leading to higher actin-myosin activity or transcription of MYH9 leading to production of non-muscle myosin (24, 40). The dimensionless parameter *λ* captures the level of nuclear-cytoskeletal crosstalk and defines the relative contribution of *ϕ_n_* to total mechanoactivation. Higher values of *λ* bias the net mechanoactivation *ϕ* towards nuclear mechanotransduction *ϕ_n_* (discussed ahead), which operates at relatively slower kinetics compared to cytoskeletal mechanotransduction. We performed parametric scans for *λ* and found that smaller values of *λ* resulted in lower mechanoactivation (*ϕ*), i.e., greater adaptation to the soft matrix, during the memory phase (Fig. S3; discussed ahead in greater detail). For *λ* = 4, the mechanoactivation trend matched well with our prior experiments of stiff-primed epithelial cells migrating on a soft matrix (24). Based on this validation, we chose *λ* = 4 for all control cases.

### Cytoskeletal mechanotransduction due to current matrix

We described the ability of cells to continually sense matrix stiffness through focal adhesion-based signaling and respond through actin-myosin force machinery in terms of a direct cytoskeletal mechanotransduction parameter *ϕ_c_*. This parameter captures the mechanisms of classic mechanotransduction due to the ability of cells to match their mechanoactivation response according the stiffness of their current adhered matrix (12, 25). Thus, we assume that *ϕ_c_* directly follows the signals from the matrix stiffness *m*, 0 < *m* ≤ 1 (Fig. 1), written as:

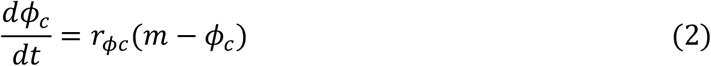

For the chosen rate constant *r*_*ϕc*_ = 6 *h*^−1^, the cytoskeletal mechanoactivation *ϕ_c_* reaches its steady state value in ~8 hours, which is consistent with our experimental observations that epithelial cells achieve stable adhesion and spreading on a given substrate in about 8 hours after seeding. According to our parametric scans of *r*_*ϕc*_, although variations in *r*_*ϕc*_ alter slopes of *ϕ* curves, the overall memory duration remains unchanged (Fig. S4), i.e., the memory acquisition and degradation is not highly sensitive to the kinetics of direct mechanotransduction.

Cytoskeletal mechanoactivation depends on matrix signal, *m*, which depends on ECM stiffness, as following:

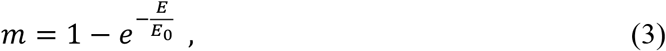

where *E* (*kPa*) is the stiffness of the underlying matrix and *m* is dimensionless. This expression is chosen based on previous work which shows that beyond a critical stiffness, focal adhesion dynamics and cell spreading are not likely to be sensitive to rise in matrix stiffness beyond a critical value (41–43). Here, *E*_0_ = 2.2 *kPa* is a calibrated matrix stiffness constant such that the matrix signal does not appreciably rise as matrix stiffness continues to increase.

### Stiffness-dependent mechanosensitive mRNA production

Cytoskeletal structure and forces are now known to regulate nuclear transport of mechanosensitive transcriptional activators such as YAP and MRTF-A (23, 44, 45). Previously, YAP has been shown to undergo phase-mediated reorganization within nucleus and cytoplasm, which leads to upregulation of YAP-related transcription factors, such as TEAD1 (46). Upregulated YAP levels have also been shown to predict higher contractility in cancer associated fibroblasts, leading to matrix stiffening and cancer cell invasion (40). In endothelial cells, persistent and steady state cell migration depends on a YAP-mediated transcriptional feedback to cytoskeletal mechanosensing (47). Additionally, microRNA-21 has been shown to enhance levels of αSMA and fibrotic mechanical memory of stem cells (22). Our previous work showed that nuclear YAP localization due to stiff-priming of epithelial cells enhances their pMLC levels and migration speed in future soft ECMs (24). Based on these established connections between mechanosensitive transcription, nuclear-cytoskeleton factor production, and overall cellular response, we assume that cellular mechanoactivation (*ϕ*) leads to mechano-sensitive transcriptional activity, which leads to production of mechanosensitive mRNA (normalized levels are denoted by *ψ*). Thus, the kinetics of mRNA levels is dependent on the cellular mechanoactivation *ϕ*, as following:

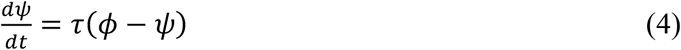

Here, mRNA levels (*ψ*) due to mechanosensitive transcription approaches cellular mechanoactivation (*ϕ*) at a rate dictated by a state of mechanosensitive epigenetic plasticity (Fig. 1). Since the epigenetic state of the cell is known to fundamentally regulate transcription factor (TF) binding and gene expression (32–34), the proposed transcription-epigenetic feedback is an important one to establish. Here, higher values of epigenetic plasticity *τ* (0, 1] are assumed to reduce chromatin resistance, enhance genome accessibility, and promote new transcription activity, thereby mRNA synthesis happens according to current stiffness. When the epigenetic plasticity is low, the DNA structure is not amenable to change and no new transcription is permitted, which leads to steady mRNA levels, determined by the previous transcriptional activity. Low epigenetic plasticity also makes it difficult to expose new TF binding sites, again disabling new transcriptional activity, thus locking the mRNA levels in place. Consistent with these observations, as described above in Eq. (4), the kinetics of mRNA levels 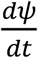 is intimately tied to epigenetic plasticity *τ*.

### Epigenetic plasticity due to priming

We model epigenetic plasticity (*τ*) in terms of cellular mechanoactivation (*ϕ*) and epigenetic modification factors (*α*) that may be generated during cell priming. Cytoskeletal structure and forces alter nuclear shape (48), which in turn regulates chromatin structure (49–53) and influences transcription of various genes (54), and in our case, rate of mRNA levels. Mechanical strain and loading have also been shown to modulate chromatin remodeling enzymes like histone deacetylase (HDAC) and histone acetyltransferase (HAT) (52) and induce rapid chromatin condensation (52). Increasing contractility has been shown to regulate levels of HDACs, whereas available spreading area has been implicated in modulating HAT (55). Recently, it has been shown that stiff-primed stem cells continue to express an epigenetic modification factor HAT1 and suppress histone deacetylation enzymes (HDACs) even after they move to a soft matrix (35). Thus, prior stiff-primed epigenetic state persists during the memory phase, which in turn could disable new chromatin remodeling and opening of TF binding sites required for soft adaptation. Matrix stiffness is also known to alter nuclear structure (27–31), which could separately influence chromatin configuration. Based on these ideas, we cumulatively define epigenetic modifiers (e.g., HAT1, HDACs) and nuclear structure regulators (e.g., Lamin A/C) in terms of ‘memory-regulating factors’ *α*, whose presence along with mechanoactivation (*ϕ*) reduces epigenetic plasticity *τ* according to chemical and mechanical switches, defined as:

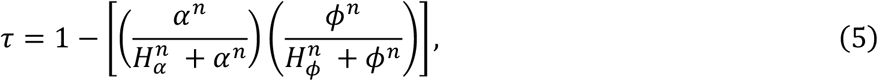

Here, 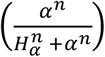 represents contribution to epigenetic plasticity due changes in genome accessibility via processes such as DNA methylation and histone modification, mediated by presence of epigenetic modification factors (*α*). The 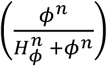 term represents changes in chromatin organization due to changes in nuclear structure caused by cellular mechanoactivation and forces. Both these terms are modeled as switches based on Michealis-Menten equations (56), where *H* is a so called half-max term that represents a point of inflection in the switch’s behavior. *H*_*α*_ and *H*_*ϕ*_ are the respective thresholds beyond which the switches are active. *H*_*α*_ = 0.5 has been chosen as the half of maximum value *α* can take. Similarly, we chose 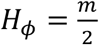, because the maximum steady state value of *ϕ* is dictated by *m*. Here, larger values of the parameter *n* (chosen here as *n* = 32) ensures that these terms, 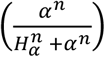 and 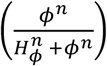, operate as binary switches that flip from ‘0’ to ‘1’ with a slight change around the critical thresholds *H*_*α*_ or *H*_*ϕ*_.

### Kinetics of memory-regulating factors

We propose that memory factors are produced only after the cell is sufficiently primed, which occurs when the mechanosensitive mRNA levels inside the cell matches the matrix signal, i.e., Δ*ψ*_*m*_ = |*m* − *ψ*| ≈ 0, defined as a binary switch of memory-specific gene expression according to:

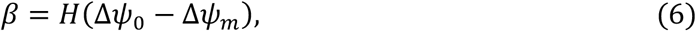

Here, *H* is the Heaviside function that outputs 1 or 0 for positive or negative inputs, respectively. The memory-specific gene expression *β* will output “1” only when the mRNA levels match the signal provided by the matrix stiffness, i.e., when Δ*ψ*_*m*_ ⟶ 0 (implemented as less than a set minimum threshold Δ*ψ*_0_ = 0.05). The kinetics of memory factors *α*_*stiff*_ and *α*_*soft*_ corresponding to stiff and soft priming, respectively, is written as:

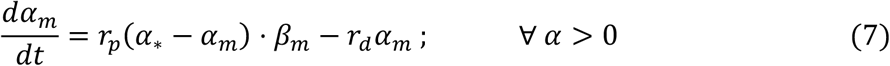

where *α*_∗_ = 1 is the maximum level of memory-regulating factors within a cell. In our previous experiments (24), 1-day priming is insufficient to generate memory response, and 3-day priming produces about 3 days of memory response, which degrades within 12-24 hours after those 3 days. According to these measurements, we calibrated the rate constants for factor production *r*_*p*_ = 0.1 *h*^−1^ and degradation *r*_*d*_ = 0.007 *h*^−1^ such that 3-day priming results in roughly 3 days of memory storage. Influence of the key parameters that regulate factor kinetics (*r*_*p*_, and *r*_*d*_) on memory regulation is presented and discussed ahead (Fig. 3).

### Nuclear contribution to net mechanoactivation

As we have noted above, epigenetic modifications due to mechanoactivation and memory regulating factors lead to maintained levels of mechanosensitive mRNA that contribute to net mechanoactivation of the cell, which ultimately causes broad cellular responses such as migration and differentiation. Indeed, previous studies have shown that upregulation of mechanosensitive transcriptional activators, such as YAP, MRTF, can proactively enhance cellular spreading, invasion, forces, growth, and migration (22, 23), which we have collectively defined as mechanoactivation. Recent work has demonstrated that YAP/TAZ regulate cytoskeletal and FA remodeling through a transcriptional feedback to enable persistent endothelial cell migration (57), highlighting the importance of transcriptional contribution to mechanoactivation. Along these lines, we define this transcriptional contribution towards cellular mechanoactivation as *ϕ_n_* (Fig. 1):

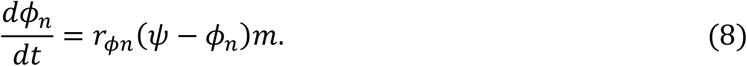

where *r*_*ϕn*_ is a rate constant that regulates how fast nuclear mechanotransduction follows the signaling due to upstream mRNA levels. Here, *ϕ_n_* represents net mechanosensitive proteins translated due to mechanosensitive mRNA transcription. Given that gene splicing followed translation and protein production are known to occur in 10-20 mins (58), we chose *r*_*ϕn*_ = 6 *h*^−1^. We performed parametric scans for *r*_*ϕn*_ and found that values of *r*_*ϕn*_ > 1 provide a sufficient feedback between transcription and mechanoactivation and higher values of *r*_*ϕn*_ do not affect mechanoactivation (Fig. S4). For a very small value of *r*_*ϕn*_ = 0.1, the protein production lags so far behind transcription that mechanoactivation and priming do not occur (Fig. S4).

Key model constants, rationale for value estimates, their parametric scans and overall effect on memory regulation are summarized in Table 1 and discussed in sections ahead.

**Table 1.**
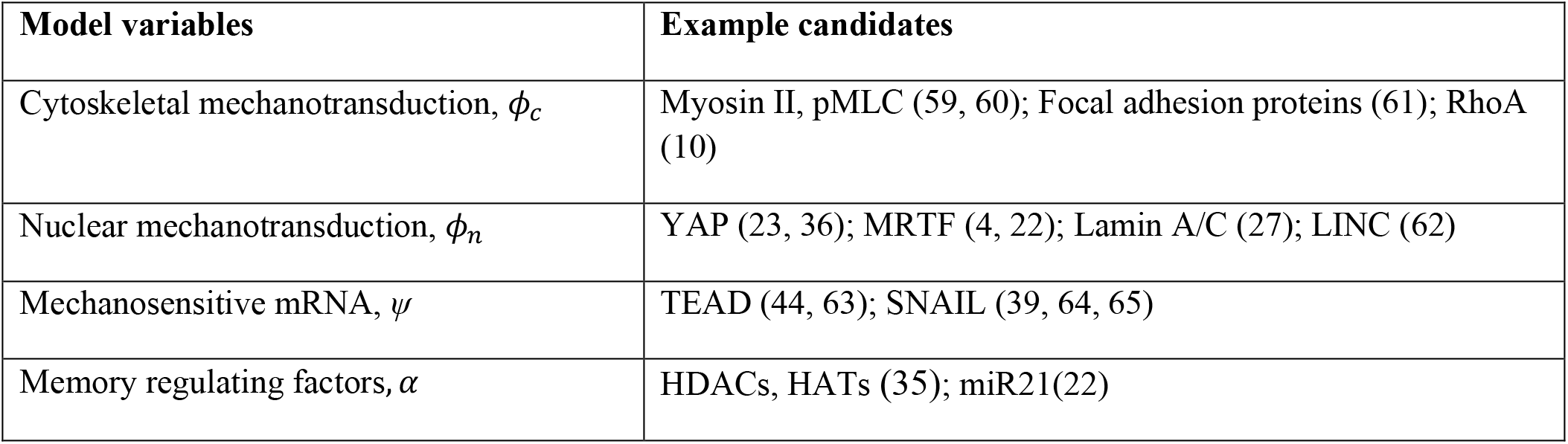
Key model variables and examples of corresponding biological candidates.

| Model variables | Example candidates |
| --- | --- |
| Cytoskeletal mechanotransduction, $\phi_c$ | Myosin II, pMLC (59, 60); Focal adhesion proteins (61); RhoA (10) |
| Nuclear mechanotransduction, $\phi_n$ | YAP (23, 36); MRTF (4, 22); Lamin A/C (27); LINC (62) |
| Mechanosensitive mRNA, $\psi$ | TEAD (44, 63); SNAIL (39, 64, 65) |
| Memory regulating factors, $\alpha$ | HDACs, HATs (35); miR21(22) |

## RESULTS & DISCUSSION

### Mechanical memory and adaptation in cells transitioning across stiff and soft matrices

Using the model described above, we calculated how the key variables corresponding to cellular mechanotransduction – mechanosensitive mRNA levels *ψ*, mechanoactivation from direct cytoskeletal signaling *ϕ_c_*, and net mechanoactivation *ϕ* – evolve overtime on matrices of defined stiffness (*m*). For a cell adhered to a matrix of Young’s Modulus *E* = 50 *kPa*, referred to as “stiff” hereafter, all signals rise and approach the matrix signal *m* = 1 (Fig. S1A). Similarly, on a matrix of *E* = 0.5 *kPa*, referred to as “soft” hereafter, signals rise to *m* = 0.2 (Fig. S1B). Both of these responses can be easily captured by existing models of mechanotransduction, in which cellular state directly follows the signals provided by the matrix stiffness.

We have previously shown that epithelial cells migrate faster on stiff (50 kPa) matrices compared to the soft (0.5 kPa) ones, and that stiff-primed cells continue to migrate fast on soft matrices (24), as qualitatively illustrated in Fig. 2A (top panel). To capture these experiments, we perform a calculation in which a cell experiences a stiff matrix for 3 days and then a soft matrix for the following 5 days – priming and migration regimen similar to experiments (24). From these model calculations, in Fig. 2A (bottom panel), we plotted net mechanoactivation *ϕ* of stiff-primed cells on soft matrix (purple line), which stays high for ~3 days before adapting to the current soft matrix. Our model calculations agree with experimental measurements of ~3 days of memory response resulting from 3 days of stiff-priming. Next, we use our model to better understand how various subcellular components operate over time through different stages of priming, memory, and adaptation.

**Figure 2:**
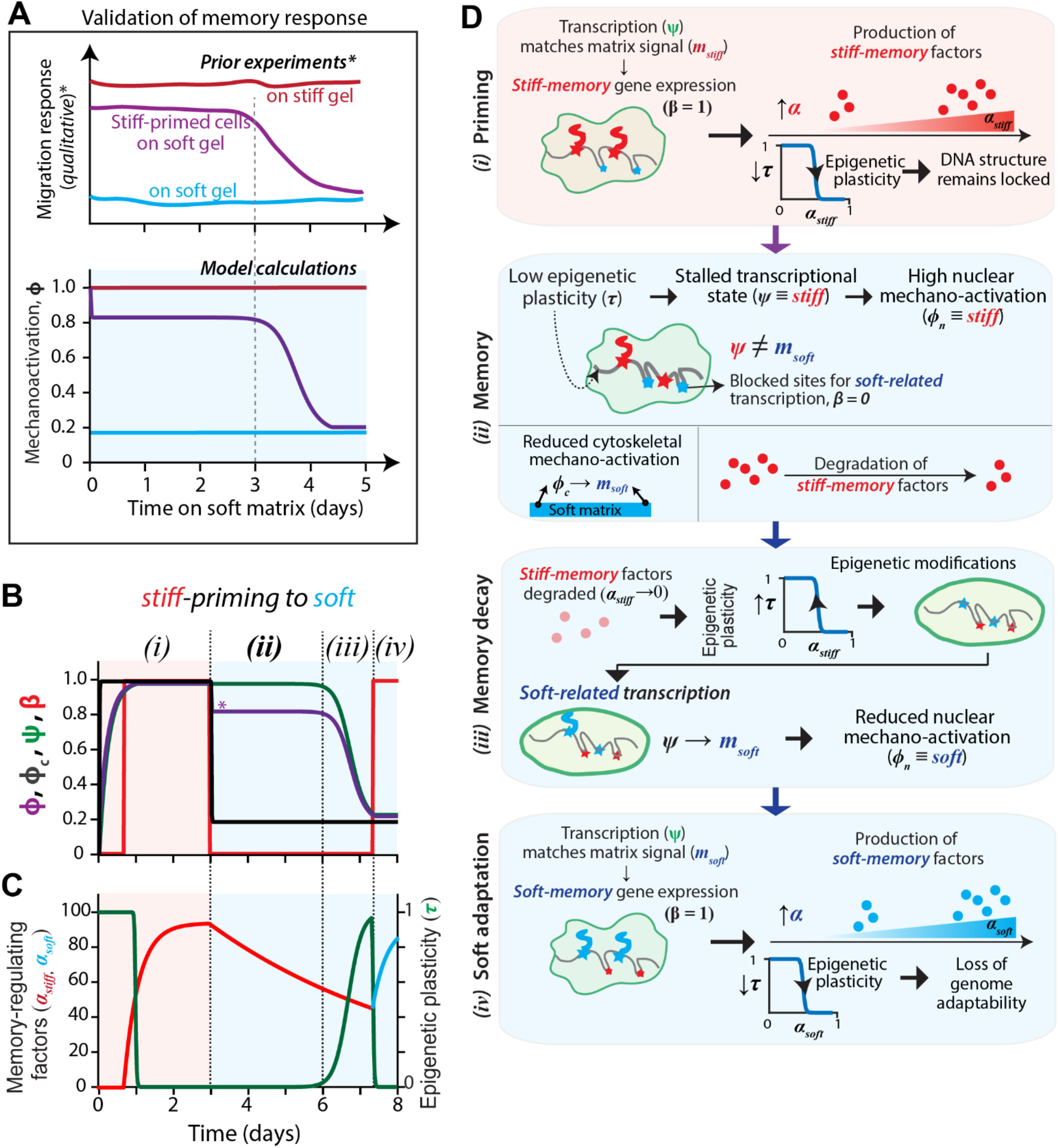
Temporal evolution of cellular mechanoactivation and transcription from stiff priming onto soft matrices. **(A)** Top panel qualitatively describes prior experimental results (24) of mechanical memory in epithelial cells primed for 3 days on stiff and soft matrices. Validation of modeling results in the bottom panel, showing mechanoactivation in cells under conditions corresponding to the experimental results – cells on soft ECM after 3-day stiff priming, compared with stiff and soft control conditions **(B)** Plots for the four key signals – transcription (*ψ*), net mechanoactivation (*ϕ*), mechanoactivation from direct cytoskeletal signaling (*ϕ_c_*), and memory-regulating transcription (*β*) – over time for a cell primed on stiff matrix (*E*=50kPa) for 3 days and then transitioning to a soft matrix (*E*=0.5kPa). In all figures, red and blue backgrounds refer to current stiff and soft matrices, respectively. * indicates plots validated against experiments by Nasrollahi *et al.* (24), in which epithelial cell migration speed showed a memory response after the 3-day stiff-or soft-priming used in these calculations. **(C)** Plots for epigenetic plasticity (*τ*) and levels of memory-regulating factors corresponding to stiff-memory (*α_stiff_*) and soft-memory (*α_soft_*). **(D)** Schematic of key steps involved in memory storage and degradation of memory for an example case of a stiff-primed cell adapting to a soft matrix, as annotated in (B): (*i*) Production of stiff-memory factors *α_stiff_* begins according to Eq. 7 when the transcription state adapts to the matrix signal, which in turn reduces epigenetic plasticity *τ*. (*ii*) When this stiff-primed cell moves to a soft matrix, new transcription does not occur immediately, because the sites for new transcription are blocked due to reduced epigenetic plasticity. This stalled transcription towards gene expressions required for adaptation to the current soft matrix means that the cellular mechanoactivation is stuck in its stiff memory, i.e., both *ψ* and *ϕ_n_* maintain their stiff levels. Here, robust priming (high *α*), low plasticity (*τ*), leads to stalled transcription, giving rise to memory. (*iii*) After the stiff-memory factors (*α_stiff_*) degrade to negligible levels, the epigenetic plasticity rises again, thus enabling epigenetic modifications, opening new TF sites and transcription for soft-adaptation commences. (*iv*) Finally, the adaptation to the new soft matrix follows the same kinetics as stiff-priming described in (*i*).

Over the first 3 days of stiff priming (Fig. 2A), mechanoactivation *ϕ* and transcription *ψ* states evolve to reach their highest level. During this priming phase, when the mechanosensitive mRNA level *ψ* matches the matrix signal m, the memory-regulating gene expression commences, i.e., *β* = 1 (Fig 2B, red line; Fig. 2D-*i*) and the level of memory-regulating factors corresponding to stiff-priming, *α*_*stiff*_, rises, which activates the epigenetic switches in Eq. 5 (Fig. 2B red line; Fig. 2D-*i*). This rise in memory-regulating factors lowers epigenetic plasticity *τ* (Fig. 2A, magenta line; Fig. 2D-*ii*).

After the cell moves to a soft matrix, the cytoskeletal mechanoactivation signal *ϕ_c_* drops quickly (Fig. 2B), which would be expected from direct mechanosensing of the current soft matrix – something a conventional mechanotransduction model would predict. However, the reduced epigenetic plasticity due to stiff-memory factors (*α*_*stiff*_) leads to a stalled transcription of genes that are necessary for adaptation to the soft matrix. This delayed adaptation to soft matrix is manifested as a memory of prior stiff priming (illustrated in Fig. 2D-*ii*). As a result, the net mechanoactivation *ϕ* reaches a second stable state of high mechanoactivation for about 3 days. Eventually, once the stiff-memory factors (*α*_*stiff*_) are degraded enough (Eq. 7) and the epigenetic plasticity *τ* increases, the new transcription for soft-specific genes commences and the cell adapts to the current matrix stiffness (Fig. 2B; illustrated in Fig. 2D-*iii*). Once the cytoskeletal signal *ϕ_c_* matches the current soft substrate, the cell now begins priming on the new substrate, and soft-primed factors (*α*_*soft*_) get produced (Fig. 2C, cyan line; Fig. 2D-*iv*). From these calculations, the plot of net mechanoactivation *ϕ* in Fig. 2B validates our prior experimental observation on epithelial cell migration speed (24), in which stiff-primed cells continued to migrate fast for approximately 3 days after arriving on to a soft matrix (Fig. 2A). With this validation in place, we next perform calculations to further understand how varying cell types defined by their ability to generate and degrade memory-regulating factors can affect memory kinetics.

### Varying kinetics of memory-regulating factors alter memory duration

During priming, cells produce memory-regulating factors and reduce their epigenetic plasticity. In previous studies, it has been shown that YAP and microRNA-21 expressions regulate short- and long-term memory responses, respectively (22–24), and the epigenetic modifiers HAT1 and HDACs may serve as memory-keepers (35), all of which could serve as memory-regulating factors. It is also likely that different cell types vary in their ability to maintain memory factors (*α*) due to varying rate constants for production (*r*_*p*_), due to translation, and degradation (*r*_*d*_), due to autophagy and protein half-life. Indeed, previous experiments (23, 35) reveal that in MSCs on stiff substrates, YAP and RUNX2 expression continues to rise over the course of 5-7 days, unlike epitheilial cells whose YAP levels reach a plateaue after 1-2 days (24), indicating varying kinetics of memory factors in these two different cell types. Here, we perform calculations to predict how cell types of varying levels and kinetics of memory-regulating factors (*α*) produce memory responses after a defined 3-day priming regimen.

For a fixed factor degradation rate constant, *r*_*d*_ = 0.1 *h*^−1^, our calculations revel that higher values of factor production rate constant *r*_*p*_ lead to higher levels of stiff-memory factors in the given 3-day priming duration (Fig. 3A). Here, *r*_*p*_ = 0.5 *h*^−1^ (Fig 3A, green line) allows the factor levels to reach their maximum value (1) during the fixed 3-day priming phase. Since, higher levels of stiff-memory factors will requires longer duration to degrade, the faster factor production rate enhances memory duration. Here, *r*_*p*_ = 0.5 *h*^−1^ generates ~3.5 days of maintained mechanoactivation (Fig 3B, green line) after 3-day priming. Very low production rate constant (*r*_*p*_ = 0.01 *h*^−1^) does not produce adequate enough memory factors, which degrade rapidly and thus resulting in no memory response (Fig 3B, cyan line). We noted that changing *r*_*p*_ does not alter memory decay phase, as all 5 curves of *α* in Fig. 3B have similar slopes in the soft matrix. For any given condition, if *α* reaches the maximum allowable level of 1, further rise in the production rate *r*_*p*_ does not yield any additional memory.

**Figure 3:**
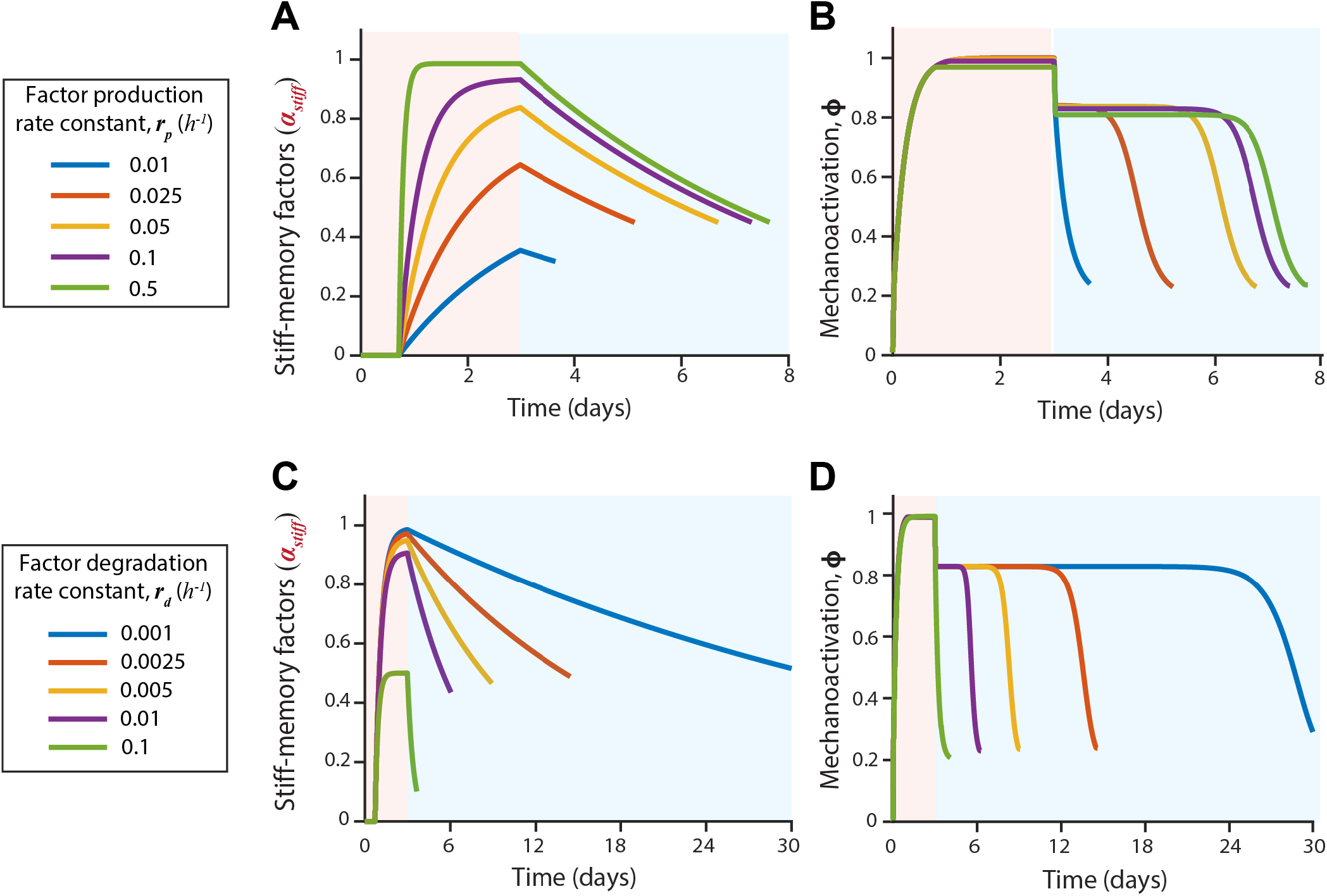
Varying kinetics of memory-regulating factors alter memory response. **(A)** Levels of stiff-memory regulating factors (*α*_*stiff*_) and **(B)** net mechanoactivation over time for cells transitioning to a soft matrix after 3 days of stiff priming, for varying rates constants of factor production *r_p_* = 0.01, 0.025, 0.05, 0.1, 0.5 *h*^−1^. **(D)** Levels of stiff-memory regulating factors (*α*_*stiff*_) and **(E)** net mechanoactivation over time for cells transitioning to a soft matrix after 3 days of stiff priming, for varying rates constants of factor degradation *r_d_* = 0.001, 0.0025, 0.005, 0.01, 0.1 *h*^−1^.

To understand how memory factor degradation affects memory response, we chose a fixed factor production rate constant *r*_*p*_ = 0.1 *h*^−1^ and plotted memory regulating factors over time for various values of *r*_*d*_. For high values of degradation rate constant *r*_*d*_ = 0.1 *h*^−1^, the simultaneous rapid degradation does not allow *α* to rise beyond 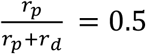, as observed in Fig. 3C, green line. As a result, no memory response is observed (Fig. 3D, green line). As the degradation rate constant *r*_*d*_ is reduced, it causes higher values of *α* at the end of the memory phase (Fig. 3C), which results in longer memory duration. For a very low degradation rate constant of *r*_*d*_ = 0.001 *h*^−1^, we predict ~21 days of memory resulting from 3 days of priming (Fig. 3D blue line).

### Longer priming yields more persistent mechanical memory

Previous experimental observations in the context of epithelial cell migration speed (24) have shown that 1-day priming is insufficient to cause memory response (qualitatively illustrated in Fig. 4A, top panel, blue line). Additionally, experiments in the context of stem cell differentiation (22, 23) have shown that longer priming, varied between 1-10 days, results in more stable memory of prior priming (qualitatively illustrated in Fig. 4A, top panel).

**Figure 4:**
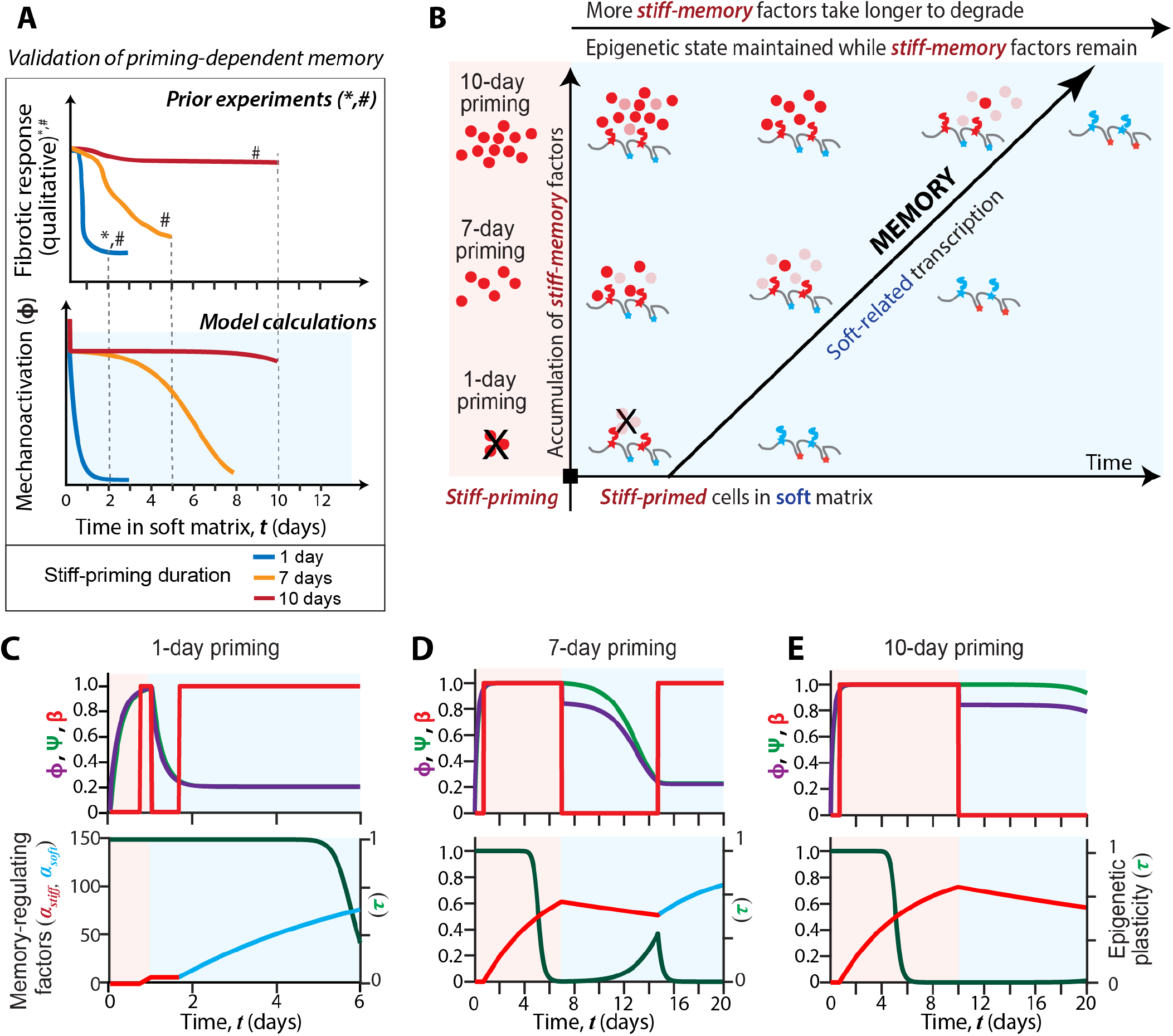
Longer priming predicts more stable memory. **(A)** Qualitative description of experiments conducted to test mechanical memory by varying priming duration from previous studies. In ^#^Yang *et al.* (23), human mesenchymal stem cells stored nuclear YAP according to past matrix stiffness after 10-day priming but not for 1-day and 7-day priming. **(A)** Bottom panel shows model predictions for mechanoactivation on soft ECM for 1-day, 7-day and 10-day priming on stiff ECM. **(B)** Schematic describing possible states of stiff-memory factors and availability of transcription factor binding sites due to epigenetic plasticity overtime for varying priming durations. Red and blue represent MRFs and binding sites corresponding to stiff and soft mechanotransduction, respectively. **(C)** 1-day, **(D)** 7-day, and **(E)**10-day priming on a stiff matrix and transferring of cells to a soft matrix, plots of net mechanoactivation level (*ϕ*), transcriptional activity (*ψ*), and stiffness specific transcription *β* in the top row and epigenetic plasticity (*τ*) and levels of memory-regulating factors corresponding to stiff-memory (*α_stiff_*) and soft-memory (*α_soft_*) in the bottom row.

To understand how our model accounts for varying priming durations, we repeated calculations with the same model constants used in the previous section. Here, cells were first stiff-primed for 1, 7 or 10 days, and subsequently allowed to reside in a soft matrix (23). We plotted net mechanoactivation *ϕ* (Fig. S2A) and found that after 1-day priming, mechanoactivation *ϕ* immediately starts to decrease as soon as it enters the soft matrix, i.e., does not result in a memory response (Fig. S2A, blue line), which is consistent with previous experiments (22–24). However, the 7-day and 10-day priming regiment both yielded ~3 days of memory (Fig. S2A orange and yellow lines), which does not match the experimental observations where 7-day priming of stem cells did not produce memory (23). This mismatch between modeling predictions and experiments could be because these calculations for stem cells use the modeling constants earlier calibrated for epithelial cells.

As we discussed in the previous section (Fig. 3), different cell types can be defined by varying factor kinetics. We then sought for find differences in epithelial and stem cells in terms of the kinetics of mechanoactivation. As illustrated in Fig. S2B, reproduced from (23) and (24), nuclear YAP localization during stiff priming rises more rapidly in epithelial cells (24) compared to stem cells (23). Thus, the factor production rate of stem cells could be slower than the epithelial cells. Next, to qualitatively understand the differences in degradation of signals in these two cell types, we illustrated how mechanoactivation decays after stiff priming (Fig. S2C), reproduced from (23) and (24). We observed that migration speed of stiff-primed epithelial cells dropped more rapidly after the memory phase (black line in Fig. S2B) compared to a slower degradation of nuclear YAP of stiff-primed stem cells after they arrived on a soft matrix. Thus, we argued that the memory factor degradation kinetics of stem cells could be slower than the epithelial cells.

Based on these qualitative differences in factor kinetics of the two cell types, we assessed that factor kinetics, both production and degradation, of stem cells are slower than the migratory epithelial cells. By utilizing the parametric scans conducted earlier (Fig. 3) and calibrating the memory response against experiments (discussed ahead), we chose *r*_*p*_ = 0.007 *h*^−1^ and *r*_*d*_ = 0.001 *h*^−1^ for stem cells. For these values, our model predicts that the 1-day and 7-day stiff-priming of stem cells yield no memory, instead the mechanoactivation *ϕ* decays steadily after arriving on the soft matrix (Fig. 4B, red line), which validates the experiments measurements (Fig. 4A). The 1-day priming regimen does not produce enough memory-regulating factors *α*_*stiff*_ to trigger the epigenetic switch (Eq. 5), i.e., the epigenetic plasticity *τ* remains high (Fig. 4C, bottom panel). As a result, the transcriptional activity does not stall and the nuclear mechanotransduction quickly adapts to new soft matrix, resulting in no memory of previous matrix stiffness. In case of 7-day priming, *α*_*stiff*_ crosses its threshold value of 0.5 and the epigenetic switch is triggered after ~5 days (Fig. 4D, bottom panel). This state of lowered epigenetic plasticity *τ* lasts only for a short duration (~ 2 days), which results in a steady decay of mechanoactivation *ϕ*, without attaining a second stable state, in the soft matrix (Fig. 4D). When we increased the duration of stiff-priming to 10 days, the mechanoactivation level remained high for ~10 days (Fig. 4E), which also validates the measured mechanical memory response in stem cells (Fig. 4A)(22, 23, 35). In this case, there is ample time for accumulation of stiff-memory factors beyond the threshold value, to ~0.9 (Fig. 4E, bottom panel). As a result, state of lowered epigenetic plasticity lasts over 10 days, resulting in a stable memory of ~10 days.

Taken together, our model calculations (Figs. 2, 4) validate experimental observations of several previous studies (22–24, 35) showing that longer priming progressively enhances the memory response, which may vary according to the kinetics of memory-regulating factors and temporal switching of epigenetic plasticity.

### Prediction of wide-ranging memory responses by tuning factor kinetics and priming duration

According to our calculations and validation against previous experiments (Figs. 2-4), our model predicts that the kinetics of memory-regulating factors define cell types (Figs. 3,4) and longer priming results in more stable memory response. Distinct combinations of these parameters couple generate the same memory response, e.g., high degradation rate can be compensated by increasing the production rate. Thus, varying kinetics of memory-regulating factors and priming durations can generate a myriad of memory responses. To better understand some of these possibilities, we calculated memory durations for a range of priming durations and the rate constants for production (*r*_*p*_) and degradation (*r*_*d*_) of memory-regulating factors (*r*_*p*_).

After 1-day stiff-priming, a case that produced no memory in our earlier calculations (Fig. 4) or any known experiments, our model predicts that increasing the factor production rate constant *r*_*p*_ > 0.1 *h*^−1^ produces a memory response, which can be further enhanced by reducing the degradation rate constant (Fig. 5A). After 3-day priming, although increasing the production rate constant *r*_*p*_ enhances the memory duration, there are diminishing returns for *r*_*p*_ > 0.08 *h*^−1^. In case of 10-day priming, the memory duration is even less sensitive to the production rate constant, where *r*_*p*_ > 0.04 *h*^−1^ do not cause appreciable change in the memory duration. Based on these plots (Fig. 5A-C), slower degradation of memory regulating factors steadily enhances memory duration as long the production rate is beyond a threshold value required to trigger the epigenetic switch discussed earlier (Eq. 5). This threshold production rate constant is lower for longer priming duration.

**Figure 5:**
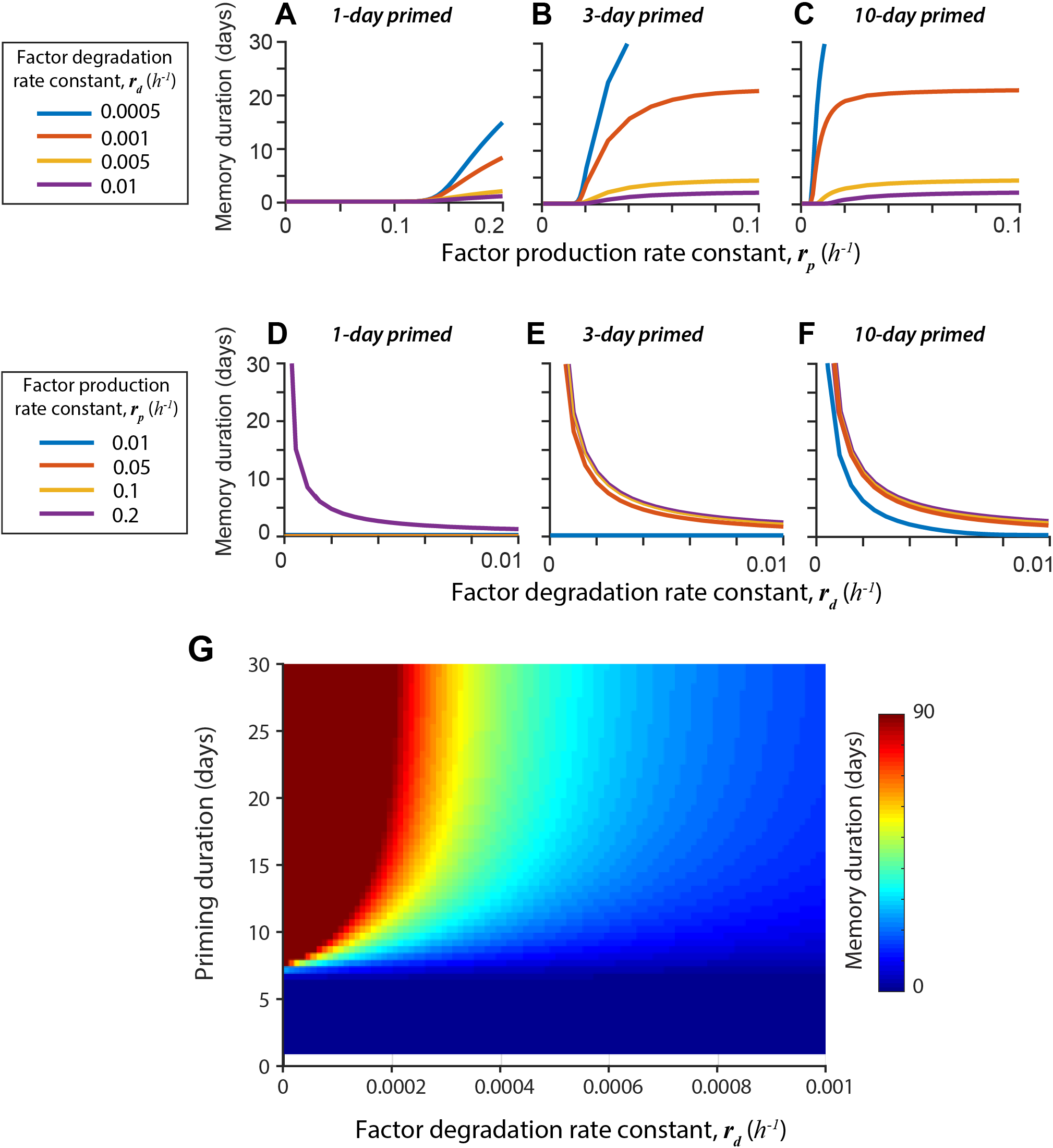
Factor kinetics and priming duration predicts wide-range of memory response. Memory duration is defined as the period (in days) over which the normalized mechanosensitive mRNA levels are within 1% of its value when it enters the soft ECM. In all simulations, cells were first primed on stiff matrix for 3 days and then transferred to a soft matrix for over 90 days. **(A-C)** Memory duration for increasing rate of production *r*_*p*_ for chosen values of factor degradation rate constant, for *(A)* 1-day, *(B)* 3-day, and *(C)* 10-day priming durations. **(D-F)** Memory duration for increasing rate of degradation *r*_*d*_ for chosen values of factor production rate constant, for *(D)* 1-day, *(E)* 3-day, and *(F)* 10-day priming durations. **(G)** Heatmap of memory duration for varying rate constants for degradation (*r*_*d*_ = 0 − 0.001 *h*^−1^) and priming duration (1-30 days), for a fixed rate constant of production *r*_*p*_ = 0.007 (corresponding to stem cells).

Next, we scanned a range of the degradation rate constant *r*_*d*_, from 0 to 0.01*h*^−1^, for four chosen values of *r*_*p*_. In case of 1-day priming, if the production rate constant *r*_*p*_ is chosen to the higher than the threshold value of 0.1 *h*^−1^, the lower values of the degradation rate constant *r*_*d*_ enhance memory duration in a hyperbolic manner, as shown for *r*_*p*_ = 0.2 *h*^−1^. Thus, even a short priming duration can be compensated by a high rate of production and low rate of degradation of the memory regulating factor. This hyperbolic dependence of memory duration on the degradation rate constant *r*_*d*_ also holds true for 3-day and 10-day priming cases, albeit with different production rate thresholds. These results show that memory duration is more sensitive to the factor degradation rate and the priming duration as compared to the factor production rate.

Finally, we picked a constant production rate constant *r*_*p*_ = 0.007 *h*^−1^ (chosen earlier for stem cells) and calculated memory duration for an even wider range of priming regimen and factor degradation rates, as shown in Fig. 5G. Here, we note that low degradation rates of memory-regulating factors can predict over 90 days of memory from less than 10 days of priming. That said, for the chosen production rate constant (*r*_*p*_ = 0.007 *h*^−1^), a minimal priming duration (~7 days)) is required to get a memory response, even in case of the slowest possible factor degradation. If the degradation rates are too high (*r*_*d*_ ≥ 0.001 *h*^−1^), even 30 days of priming does not result in appreciable memory. In sum, our model proposes that both factor kinetics and priming duration can independently regulate the memory response. Thus, we predict that shorter duration of mechanical priming could be compensated by altering the kinetics of memory-regulation factors, such as faster production and slower degradation, through precise molecular perturbations.

## CONCLUSIONS & OUTLOOK

Our model of cellular mechanotransduction builds on several established ideas of cellular sensing and response to matrix stiffness. The adaptation of cells to matrix mechanics is conventionally understood through focal adhesion dynamics and intracellular forces generated by actin polymerization and myosin-based contractility, often referred to as clutches and motors (13). In the interest of simplicity, we chose not to explicitly model those aspects and instead provided a phenological description of matching cytoskeletal and matrix signals. The presented mathematical framework focuses on upstream mechanosensitive mRNA production and epigenetic plasticity, which result in the observed cytoskeletal signaling. One of the key challenges in capturing both long-term memory of past environments and short-term adaptation to the current environment of cells is that the net cellular mechanoactivation state is a combination of processes that occur at very different timescales – seconds-minutes for focal adhesion dynamics and hours-days for transcriptional kinetics and epigenetic modifications. Here, we address this problem by expanding the framework of mechanotransduction through a feedback between cytoskeletal signaling, mechanosensitive transcription, and epigenetic plasticity (Fig. 5).

### Validation of the model against existing experimental data

Over the past decade, the nuclear-cytosolic translocation of YAP, a transcriptional co-activator of the Hippo pathway, has been recognized as a key regulator of cellular mechanosensing (36). Although the nuclear YAP localization on stiff matrices is widely established, recent studies showed that stiff-primed human mesenchymal stem cells and mammary epithelial cells continue to store nuclear YAP on soft matrices (23, 24). Conventional models for mechanotransduction would predict that cells should quickly dissociate their adhesions and weaken cytoskeletal forces on the new soft environment, and thus reduce nuclear YAP. Since our model couples the cytoskeletal signaling with the mechanosensitive transcriptional state of the cell, we are able to predict a mechanical memory in which transcriptional activity and mechano-activation are delayed upon transfer to a new environment (Fig. 2A). One important condition for this delay is sufficient duration of priming, which we captured through memory-regulating factor production along with switches in epigenetic plasticity. Consistent with the experiments (22–24), we also predict that longer priming causes more robust memory response, because we propose that the cells do not commence new transcription required for adaptation to a new environment, until the memory-regulating factors are degraded and the epigenetic plasticity is reset (Fig. 4A). In Tables 1 and 2, we summarize some example molecular candidates corresponding to each model component and the range of memory responses produced by various model parameters. Although our model captures these existing outcomes of mechanical memory in stem cells and epithelial cell migration (22–24, 35), key model components rely on the proposed biological processes (Figs. 1, 6) that will need to be experimentally validated in the future.

**Table 2.** List of rate parameters, range estimation, and resulting memory response.

| Parameter | Control value | Rationale | Range | Memory response (after 3-day priming) | Results |
| --- | --- | --- | --- | --- | --- |
| $r_{\phi_c}$, Rate of cytoskeletal mechanotransduction | 6 h<sup>-1</sup> | Focal adhesion sensing and clustering is of the order of ~10 minutes | 0.1-100 h<sup>-1</sup> | ~3 days | Fig. S4A |
| $r_{\phi_n}$, Rate of nuclear mechanotransduction | 6 h<sup>-1</sup> | Gene splicing and protein translation is of the order of ~10 minutes | 0.1-100 h<sup>-1</sup> | 0 to 3 days | Fig. S4B |
| $\lambda$, Ratio of nuclear to cytoskeletal contribution | 4 a.u. | Calibration with memory response | 1-10 a.u. | ~3 days | Fig. S3 |
| $r_p$, Factor production rate constant | 0.1 h<sup>-1</sup> | 3-day priming giving 3-day memory, which degrades in about 12 hours once memory starts to degrade. | 0-0.5 h<sup>-1</sup> | 0 to 3.5 days | Fig. 3 |
| $r_d$, Factor degradation rate constant | 0.007 h<sup>-1</sup> | Same as above | 0-0.1 h<sup>-1</sup> | ∞ to ~0 days | Fig. 3 |

(a.u. = arbitrary units)

**Figure 6:**
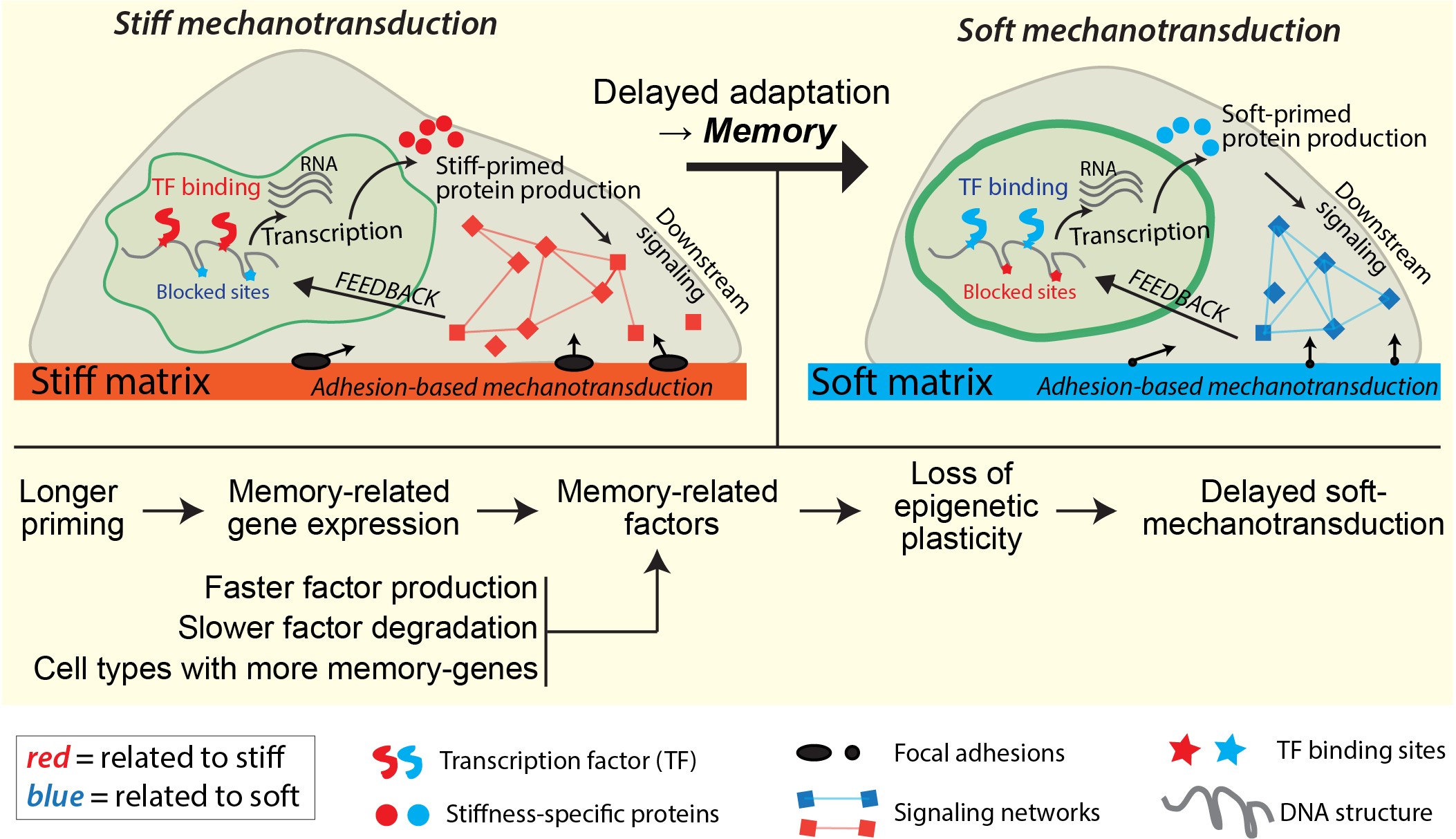
An expanded framework for mechanotransduction, including mechanical memory due to feedbacks from cytoskeletal signaling, transcription, and epigenetic plasticity. A schematic summarizing how various components of mechanotransductive detailed in this study are connected to one another for cells on stiff and soft matrices. Mechanotransduction starts with cytoskeletal signaling from the respective matrix, causing cytoskeletal mechanoactivation that affects DNA structure, availability of transcription factor binding sites, gene expression and feedback to the net cellular mechanoactivation. Red and blue color themes correspond to events and processes related to stiff and soft matrices. Based on our calculations, validation with existing experimental results, and new prediction, the key regulators of memory storage are priming duration and the state of memory-regulating factors.

### Proposed studies to test key assumptions and predictions

In the process of validating the core components of the model against prior experimental data (Figs. 2,4), we also made several predictions that can be tested in future experiments. For example, different cell types are expected to follow different kinetics of priming – a possibility we capture by varying memory factor kinetics (Fig. 3,S2). By observing accumulation and degradation of mechanosensitive markers like YAP and RUNX2, we can predict the timescales of the memory factor production and degradation. According to our predictions, cells with faster production and slower degradation rates of memory-regulating factors would generate longer memory responses, which is validated by recent experiments (35). In particular, we found that slowing the factor degradation rate had the highest advantage for the memory response, which may be experimentally tested by treating the cells with specific autophagy inhibitors and measuring memory response in different cell behaviors, such as migration and fate, as applicable according to the cell type and matrix context. We also proposed that different cell types vary in their steady state value of memory-regulating factors, which can be tested by performing proteomic analyses of different cell types after stiff-priming and measuring the number and levels of proteins and enzymes that feature in memory regulation. This information can then be used to calibrate the rate constants and predict relative memory responses of varying cell types.

Although our model integrates known biological processes, there are some key assumptions that lack direct experimental evidence at this stage. We assumed that longer priming leads to progressively higher production of memory-regulating factors and reduced epigenetic plasticity. Recent experiments support this notion by showing upregulated HAT1 and downregulated HDACs when cells are primed sufficiently (35). Although it is known that nuclear shape, chromatin structure, and epigenetic state of the cell depend on matrix stiffness (27, 30, 31, 55), it remains unclear how quickly these nuclear markers change when cells move to a new environment. Experiments could be also performed in which nuclei of differentially-primed cells are imaged through live-cell microscopy over time to verify that the nuclear structure and shape remain locked according to their past primed state. Separately, RNA-sequencing and proteomics over time could provide more specific kinetics for another assumption of our model that the memory-regulating gene expressions and factor levels vary over time, which then need to be degraded before adaptation to new environments can occur. It should be noted that the timescales for these processes might vary according to cell types and the range of matrix stiffness. We anticipate that experimental measurements across cell types and matrix contexts will help validate our model and calibrate associated parameters.

### Limitations and future work

Despite the expansive reach of our model across multiple matrix stiffnesses and scales of subcellular processes, connecting cytoskeletal signaling to upstream transcription and feedback with epigenetic plasticity, it does not explicitly model adhesion dynamics, cytoskeletal forces, and nuclear structure. As a result, our predictions do not address how various physical components within the cell evolve for varying values of matrix stiffness and over time. Overall, the model is somewhat descriptive and phenomenological in nature. However, these choices had to be made because the feedbacks among cytoskeletal signaling, memory-regulating factors, and epigenetic plasticity are currently unknown. Moreover, our goal was to focus on the core cellular mechanoactivation kinetics in the context of memory and adaptation, which remain absent in existing models for mechanotransduction. Despite its limitations, our mathematical description of mechanotransduction is the first one to capture and validate mechanical memory observed in several matrix conditions, priming regimen, and cell types. We hope that our predictions and proposed governing biological processes will motivate new experimental measurements, as noted above, which will help further tease apart the subcellular mechanisms of cellular memory and adaptation that are fundamental to cell mechanobiology.

## ACKNOWLEDGEMENTS

This work was in part supported by the NIH/NIGMS (R35 GM128764) and Siteman Cancer Center grants to AP, and the NSF Center for Engineering MechanoBiology funding to VS and AP.

## AUTHOR CONTRIBUTIONS

AP conceived and supervised the project. JM, VS, and AP formulated mathematical models. JM performed computational modeling and plotted calculations. JM, VS, and AP interpreted findings, made figures, and wrote the manuscript. VS and AP acquired funding.

## SUPPLEMENTARY INFORMATION

**Figure S1:**
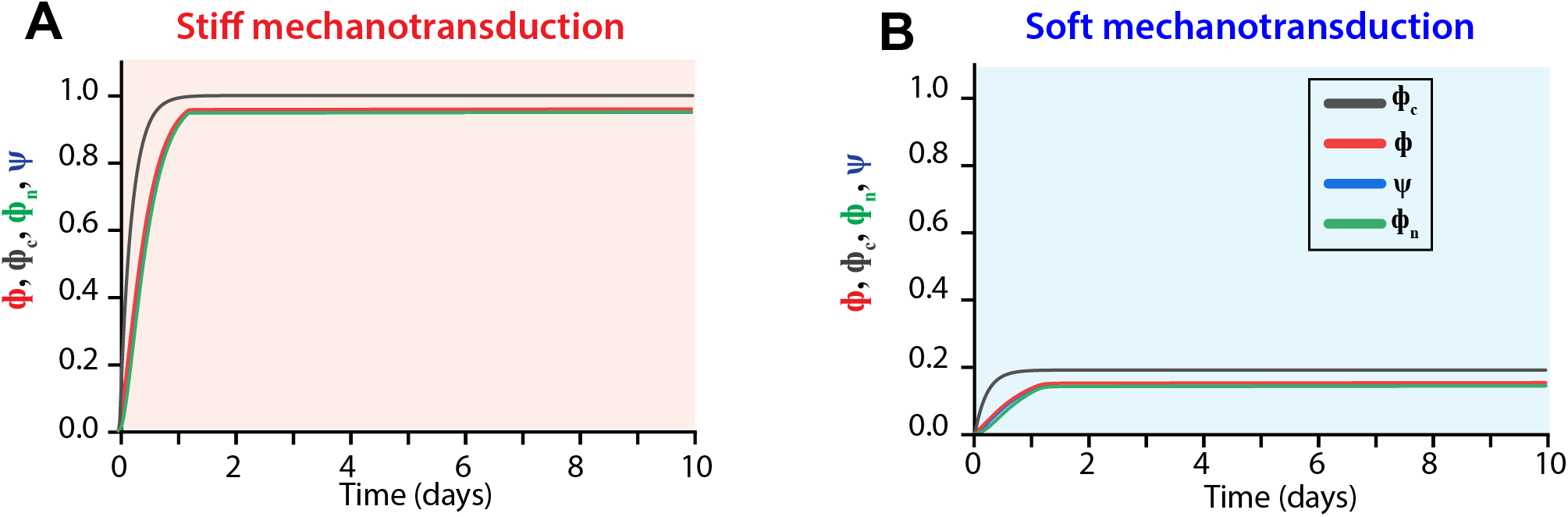
Temporal evolution of cellular mechanoactivation and transcription on stiff and soft matrices. Plots for the four key signals ‒transcriptional activity (*ψ*), mechanoactivation from the nucleus (*ϕ_n_*), mechanoactivation from direct cytoskeletal signaling (*ϕ_c_*), and net mechanoactivation (*ϕ*) – over time for control **(A)** stiff and **(B)** soft matrices.

**Figure S2:**
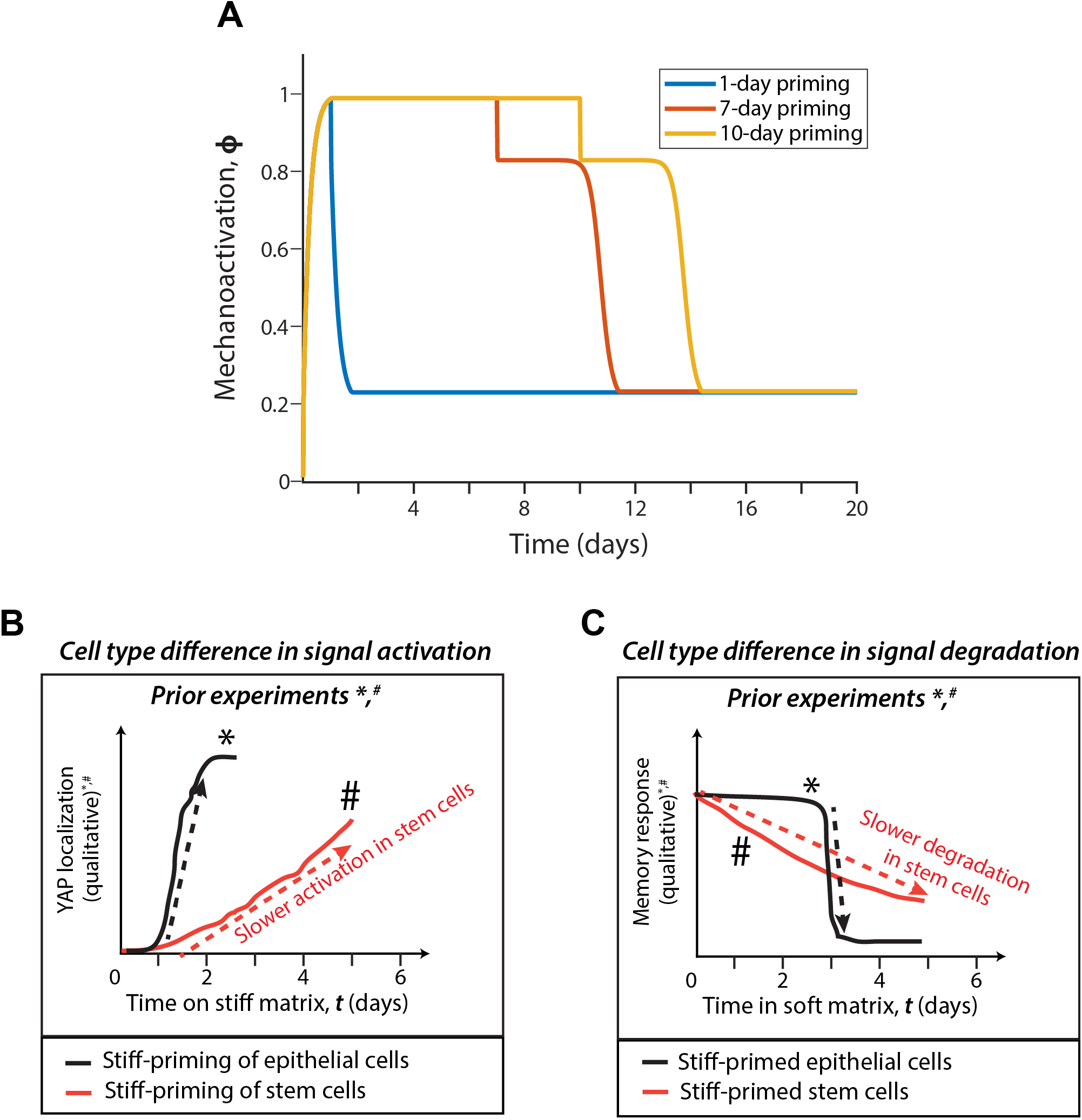
Calibrating differences between epithelial and stem cells. **(A)** Model predictions for mechanoactivation when cells stiff-primed for 1-day, 7-day and 10-day transition to the soft ECM. (B) and (C) describe qualitative description of experiments conducted to test mechanical memory by varying priming durations in two previous studies. **(B)** In *Nasrollahi *et al.* (24), epithelial cells accumulate nuclear YAP quicker, in about 2 days (black line). In ^#^Yang *et al.* (23), human mesenchymal stem cells stored nuclear YAP in about 5 days, much longer than epithelial cells (red line). **(C)** In *Nasrollahi *et al.* (24), epithelial cell migration speed showed a 3-day memory response after 3-day priming (black line), which decays within 12 hours. In ^#^Yang *et al.* (23), human mesenchymal stem cells stored nuclear YAP according to past matrix stiffness after 10-day priming but not for 7-day priming and the signal decays in about 5 days (red line).

**Figure S3:**
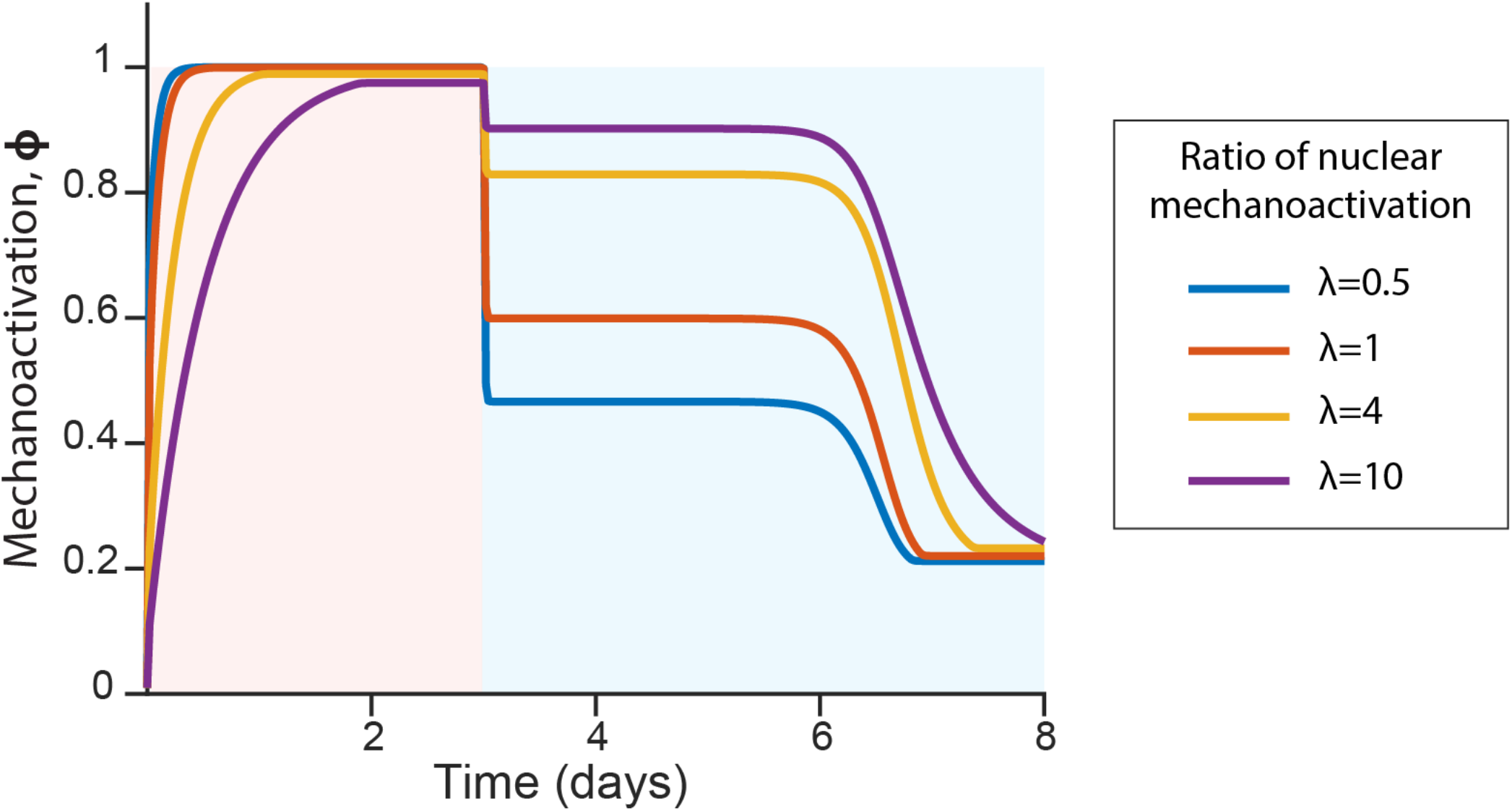
Effect of the ratio of nuclear to cytoskeletal mechanoactivation (*λ*) on mechanical memory. Calculations for net mechanoactivation (*ϕ*) over time for 3-day stiff primed cells moving to soft ECM are plotted, for varying values of the ratio (dimensionless) of nuclear to cytoskeletal mechanoactivation, *λ* = 0.5, 1, 4, 10.

**Figure S4:**
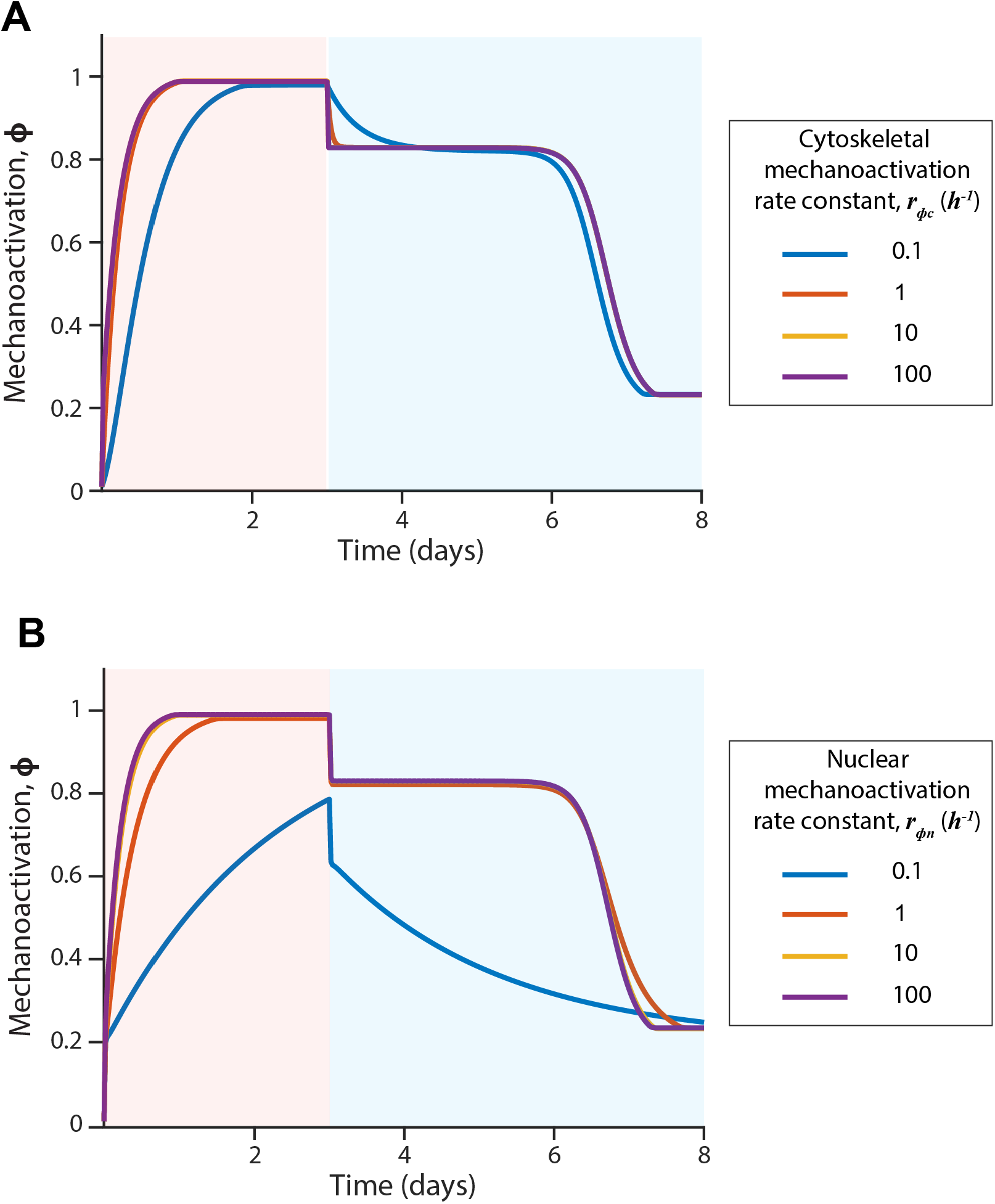
Effects of varying mechanoactivation rate constants on mechanical memory for given 3-day priming. Equation (1) states that mechanoactivation has 2 components, cytoskeletal mechanoactivation and nuclear mechanoactivation. Here, we vary their rate constants *r*_*ϕc*_ and *r*_*ϕn*_, and plot mechanoactivation over time for cells stiff-primed for 3 days and then transitioning to a soft ECM. For varying values of **(A)** *r*_*ϕc*_ = 0.1 − 100 *h*^−1^ and **(B)** *r*_*ϕn*_ = 0.1 − 100 *h*^−1^ net mechanoactivation is plotted over time.

